# The SUMO conjugation complex self-assembles into nuclear bodies independent of SIZ1 and COP1

**DOI:** 10.1101/393272

**Authors:** Magdalena J. Mazur, Mark Kwaaitaal, Manuel Arroyo Mateos, Francesca Maio, Ramachandra K. Kini, Marcel Prins, Harrold A. van den Burg

**Author notes:** Authors contributed equally to this work (Magdalena J. Mazur, Mark Kwaaitaal). Corresponding Author: Dr. Harrold A. van den Burg, Molecular Plant Pathology, SILS, University of Amsterdam, Science Park 904, 1098XH, Amsterdam, the Netherlands.

## Abstract

**One sentence Summary:** SUMO conjugation activity causes formation of SUMO nuclear bodies, which strongly overlap with COP1 bodies thanks to a substrate-binding (VP) motif in the E3 ligase SIZ1 that acts as bridge protein.

**Abstract:** Attachment of the small ubiquitin-like modifier SUMO to substrate proteins modulates their turnover, activity or interaction partners. An unresolved question is how this SUMO conjugation activity concentrates the enzymes involved and the substrates into uncharacterized nuclear bodies (NBs). We here define the requirements for the formation of SUMO NBs and for their subsequent co-localisation with the master regulator of growth, the E3 ubiquitin ligase COP1. COP1 activity results in degradation of transcription factors, which primes the transcriptional response that underlies elongation growth induced by night-time and high ambient temperatures (skoto- and thermomorphogenesis, respectively). SUMO conjugation activity itself is sufficient to target the SUMO machinery into NBs. Co-localization of these bodies with COP1 requires besides SUMO conjugation activity, a SUMO acceptor site in COP1 and the SUMO E3 ligase SIZ1. We find that SIZ1 docks in the substrate-binding pocket of COP1 via two VP motifs - a known peptide motif of COP1 substrates. The data reveal that SIZ1 physically connects COP1 and SUMO conjugation activity in the same NBs that can also contain the blue-light receptors CRY1 and CRY2. Our findings thus suggest that sumoylation apparently coordinates COP1 activity inside these NBs; a mechanism that potentially explains how SIZ1 and SUMO both control the timing and amplitude of the high-temperature growth response. The strong co-localization of COP1 and SUMO in these NBs might also explain why many COP1 substrates are sumoylated.

**Funding information:** The Netherlands Scientific Organisation (ALW-VIDI grant 864.10.004 to HvdB) and the Topsector T&U program Better Plants for Demands (grant 1409-036 to HvdB), including the partnering breeding companies, supported this work; FM is financially supported by Keygene N.V. (The Netherlands).

## Introduction

The post-translational modification of proteins by SUMO1 and -2 (SMALL UBIQUITIN-LIKE MODIFIER 1 and 2) is an essential process in Arabidopsis (*Arabidopsis thaliana*) affecting hundreds of proteins (Saracco et al., 2007; Miller et al., 2010; Rytz et al., 2018). These two closely related proteins are the nearest homologs of the ancestral SUMO protein (Hammoudi et al., 2016). Their attachment to substrates is catalysed in two steps by the SUMO E1 ACTIVATING ENZYME (SAE1/2) dimer and the SUMO E2 CONJUGATING ENZYME (SCE1) (Colby et al., 2006; Saracco et al., 2007). SCE1 directly recognizes and conjugates a SUMO moiety to the acceptor lysine embedded in a short consensus motif ψKxE (where ψ denotes a bulky hydrophobic residue, K the acceptor lysine, x any residue, and E is Glutamate) (Bernier-Villamor et al., 2002; Yunus and Lima, 2006). SUMO conjugation (sumoylation) can also be promoted by SUMO E3 ligases, but the conditions by which E3s are recruited to SCE1 remain unclear (Gareau and Lima, 2010). SUMO proteome analyses with human cells suggested that −50% of the SUMO targets are modified at non-consensus sites (Hendriks et al., 2014; Lamoliatte et al., 2017). Structural studies exposed that the E3 ligases orientate the E2 enzyme such that they favour SUMO donor transfer to non-consensus lysines (Yunus and Lima, 2009; Streich and Lima, 2016). Alternatively, substrate selection can involve non-covalent interactions between SUMO and a short hydrophobic peptide in substrates, called the SUMO-interaction motif (SIM) (Zhu et al., 2008; Flotho and Melchior, 2013). SIM peptides bind to SUMO by forming an additional alien β-strand in the β-sheet of SUMO (Song et al., 2005; Hecker et al., 2006; Sekiyama et al., 2008).

SUMO often acts as a monomeric adduct, but it can also form SUMO chains (Colby et al., 2006). In yeast and mammals SUMO chains are important signals in meiosis, genome maintenance and (proteotoxic) stress (Vertegaal, 2007; Ulrich, 2008; Bruderer et al., 2011; Tatham et al., 2011). These SUMO chains are recognized by SUMO-targeted Ubiquitin E3 ligases (StUbls) that mark the SUMO-modified proteins for degradation (Perry et al., 2008; Guo et al., 2014). Distant homologues of these StUbls were identified in the Arabidopsis genome, but their biological function remains to be determined (Elrouby et al., 2013). In Arabidopsis, two SP-RING domain-containing proteins, PROTEIN INHIBITOR OF ACTIVATED STAT LIKE1 (PIAL1) and PIAL2, were shown to promote SUMO chain formation (Tomanov et al., 2014). There is also a role for the SUMO E2 enzyme in SUMO chain formation (Tomanov et al., 2018). The SUMO E2 interacts via two sites with SUMO: (i) a thioester bond (∼) between the catalytic cysteine in the E2 and the C-terminal glycine of SUMO (covalent SUMO loading in the catalytic pocket, E2∼SUMO) and (ii) a strong nearly constitutive, non-covalent ‘SIM-like’ interaction (non-covalent SUMO binding, SUMO·E2) (Bencsath et al., 2002; Knipscheer et al., 2007; Streich and Lima, 2016). This second non-covalent interaction between SUMO and SCE1 is apparently needed for chain formation, as mutations in the human SUMO E2 enzyme (Ubc9) that disrupt this interaction also strongly reduce SUMO chain formation *in vitro* (Knipscheer et al., 2007).

Many studies on the SUMO (de)conjugation machinery revealed that the SUMO enzymes and their targets often reside in uncharacterized punctate nuclear structures of various shapes and sizes (Murtas et al., 2003; Conti et al., 2008; Cheong et al., 2009; Castaño-Miquel et al., 2013; Xiong and Wang, 2013). The E3 ubiquitin ligase CONSTITUTIVE PHOTOMORPHOGENIC 1 (COP1), which is a SUMO substrate, also localizes to nuclear bodies (NBs) (Kim et al., 2016; Lin et al., 2016). COP1 is essential for the dark and high-temperature induced growth responses (skoto-and thermomorphogenesis, respectively) (Lau and Deng, 2012; Park et al., 2017; Hammoudi et al., 2018). Both conditions cause COP1 to translocate from the cytosol to the nucleus where it localizes to a subclass of NBs (Stacey et al., 1999; Stacey and von Arnim, 1999; Park et al., 2017), which are proposed sites for degradation of COP1 substrates (Van Buskirk et al., 2012). COP1 activity is stimulated by the SUMO E3 ligase SIZ1 and correspondingly both skoto- and thermomorphogenesis are strongly compromised in the SIZ1 null mutant *siz1-2* and the knockdown mutant *sumo1-1;amiR-SUMO2* (Delker et al., 2014; Lin et al., 2016; Park et al., 2017; Hammoudi et al., 2018). Correspondingly, we previously showed that the high temperature induced transcriptional response of many PIF4 and BZR1 genomic targets was both delayed in timing and reduced in amplitude in the *siz1-2* and *sumo1-1;amiR-SUMO2* backgrounds (Hammoudi et al., 2018). In particular, PIF4 is the master regulator of thermomorphogenesis, whose gene expression is tightly controlled by COP1 activity (Koini et al., 2009; Quint et al., 2016). Notably, COP1 contains a single SUMO acceptor site (Lys193) that is embedded in a non-consensus site and whose sumoylation strictly depends on SIZ1 (Lin et al., 2016).

As the formation and role of these SUMO and COP1 NBs is poorly understood, we examined how the SUMO conjugation complex and COP1 physically interact in NBs. We find that the formation of SUMO·SCE1 NBs is dynamic and depends on catalytic activity of the SUMO E1 and E2 enzymes. Likewise, only the conjugation-competent SUMO^GG^ form of SUMO1 can stimulate formation of the SUMO·SCE1 (SUMO·E2) and SUMO·SIZ1 (SUMO·E3) NBs, while the non-covalent SUMO·SCE1 interaction via the SIM has apparently a dual role in their formation. Co-localization of these SUMO·SCE1/SIZ1 NBs with COP1 depends on the SUMO acceptor site in COP1. Conversely, we expose that SIZ1 is a COP1-dependent ubiquitination substrate due to two VP motifs that can directly bind to the COP1 substrate binding pocket. Our data thus provide a mechanistic link between the subcellular localization of the SUMO conjugation complex and COP1 in one and the same NBs, but recruitment to these bodies depends on the intrinsic properties of the proteins involved, i.e. conjugation activity of SCE1 and substrate selection of COP1.

## Results

### Arabidopsis SUMO1 interacts with SCE1 and SIZ1 via its SIM-binding site

To this point it remains undefined whether Arabidopsis SUMO1 and -2 interact with partnering proteins via SIMs. Based on sequence homology between Arabidopsis and the human SUMOs, we mutated two conserved hydrophobic residues (Phe32, Ile34) in the β 2-strand of Arabidopsis SUMO1 to test in the Y2H assay if – in analogy to the yeast and mammalian systems – these two residues determine binding of SIM-containing proteins (Supplemental Fig. S1A). Wild type SUMO1 and the F32A+I34A mutant (Supplemental Table S1, SUMO1^*SIM*^) were expressed as bait fusions with the GAL4 binding domain (BD) and four human proteins with a known SIM were used as preys (as GAL4 AD fusions) (Hecker et al., 2006). To only test for non-covalent interactions between SUMO1 and these SIM-containing proteins, we expressed a conjugation-deficient SUMO1 variant; this variant lacks the C-terminal diGly motif needed for SUMO attachment to acceptor lysines (SUMO1*^ΔGG^*). As expected, SUMO1*^ΔGG^* interacted with the three of the four tested SIM-containing proteins and these interactions were suppressed by the F32A+I34A mutations for all except PIAS1 (Supplemental Fig. S1A; SUMO1*^ΔGG+SIM^*). Deletion of the diGly motif (‘ΔGG’) and/or introduction of the F32A+I34A double mutation (‘SIM’) did not reduce the protein accumulation of SUMO1 in yeast (Supplemental Fig. S1G). Thus, the β2-strand of Arabidopsis SUMO1 apparently facilitates binding of SIM peptides, similar to its human and yeast counterparts.

To assess if Arabidopsis SUMO1 interacts via this SIM interface with SCE1 or SIZ1, the BD-SUMO1 fusions were expressed together with SCE1 or SIZ1 fused to the GAL4 AD in the Y2H. Both SCE1 and SIZ1 interacted with SUMO1^GG^ and SUMO1^Δ*GG*^. The SUMO1-SCE1 interaction was impaired when the SIM-binding pocket was mutated (Supplemental Fig. S1, B and C, GG+*SIM* and *ΔGG*+*SIM*). The interaction between SUMO1 and SIZ1, was reduced for all SUMO1 mutant versions (Supplemental Fig. S1, B and C, *ΔGG,* GG+*SIM* and *ΔGG*+*SIM*). As the different SUMO1 variants all accumulated at least to same protein level as wild type SUMO1 in yeast (Supplemental Fig. S1G), we conclude that SIM(-like) interactions also play a role in the SUMO-SCE1 interaction. Additional support comes from our recent work where we demonstrated that a conserved SIM-like motif in the N-terminal tail of SCE1 is essential for SUMO1 binding (Mazur et al., 2017). Our findings here suggest the same for SUMO-SIZ1 protein complexes in Arabidopsis, as previously reported for the homologous proteins from yeast and mammals (Cheong et al., 2010; Mascle et al., 2013; Kaur et al., 2017); however, functional mutants of the SIM domain in SIZ1 still need to be tested.

### SUMO1 conjugation activity causes SCE1 and SIZ1 to re-localize to nuclear bodies

We then examined if these SIM(-like) interactions affected the subcellular localization of SCE1 and SIZ1 *in planta*. We first analysed the localization of the individual proteins expressing them as GFP fusions in *N. benthamiana*. YFP-SUMO1, GFP-SCE1, and GFP-SIZ1 localized to the nucleus, cytosolic pockets near the FM4-64-marked plasma membrane and in cytoplasmic strands (Supplemental Fig. S2, A to C). GFP-SUMO1 accumulated in the cytoplasm and nucleus independent of its diGly motif and SIM-binding site (Supplemental Fig. S1D). Likewise, SCE1 resided both in the cytoplasm and nucleus (Supplemental Fig. S2, B and D), while SIZ1 accumulated primarily in the nucleus, but a residual signal remained visible in the cytoplasm (Supplemental Fig. S2C).

To determine if the interaction with SUMO changes the subcellular localization of SCE1 or the E3 ligase SIZ1, the proteins were expressed as BiFC pairs. To this end SUMO1 was fused via its N-terminus to super-CFP^N^ (fragment SCFP^1-173^), while SCE1 was fused at its N-terminus to SCFP^C^ (fragment SCFP^156-239^). SIZ1 was fused at its C-terminus to SCFP^C^. Expression of both SCFP^N^-tagged SUMO1^GG^ and SUMO1^GG+*SIM*^ with SCFP^C^-tagged SCE1 or SIZ1 yielded in all four combinations CFP reconstitution in relatively large punctate structures that were exclusively found in the nucleus (as marked with Hoechst dye; Supplemental Figure S3A.2), hereafter called nuclear bodies (NBs) (Fig. 1, A and B; Supplemental Fig. S3). These NBs were absent for the BiFC combinations with SUMO1*ΔGG* or SUMO1*^ΔGG+SIM^*; instead in combination with SCE1 their BiFC signal displayed a uniform distribution in both the nucleus and cytoplasm (Supplemental Fig. S3A’), while in combination with SIZ1 their signal was homogeneously spread only in the nucleus (Supplemental Fig. S3B’). These findings suggest that SUMO conjugation itself causes NB assembly in the case of the SUMO1·SCE1 and SUMO1·SIZ1 BiFC pairs.

**Figure 1:**
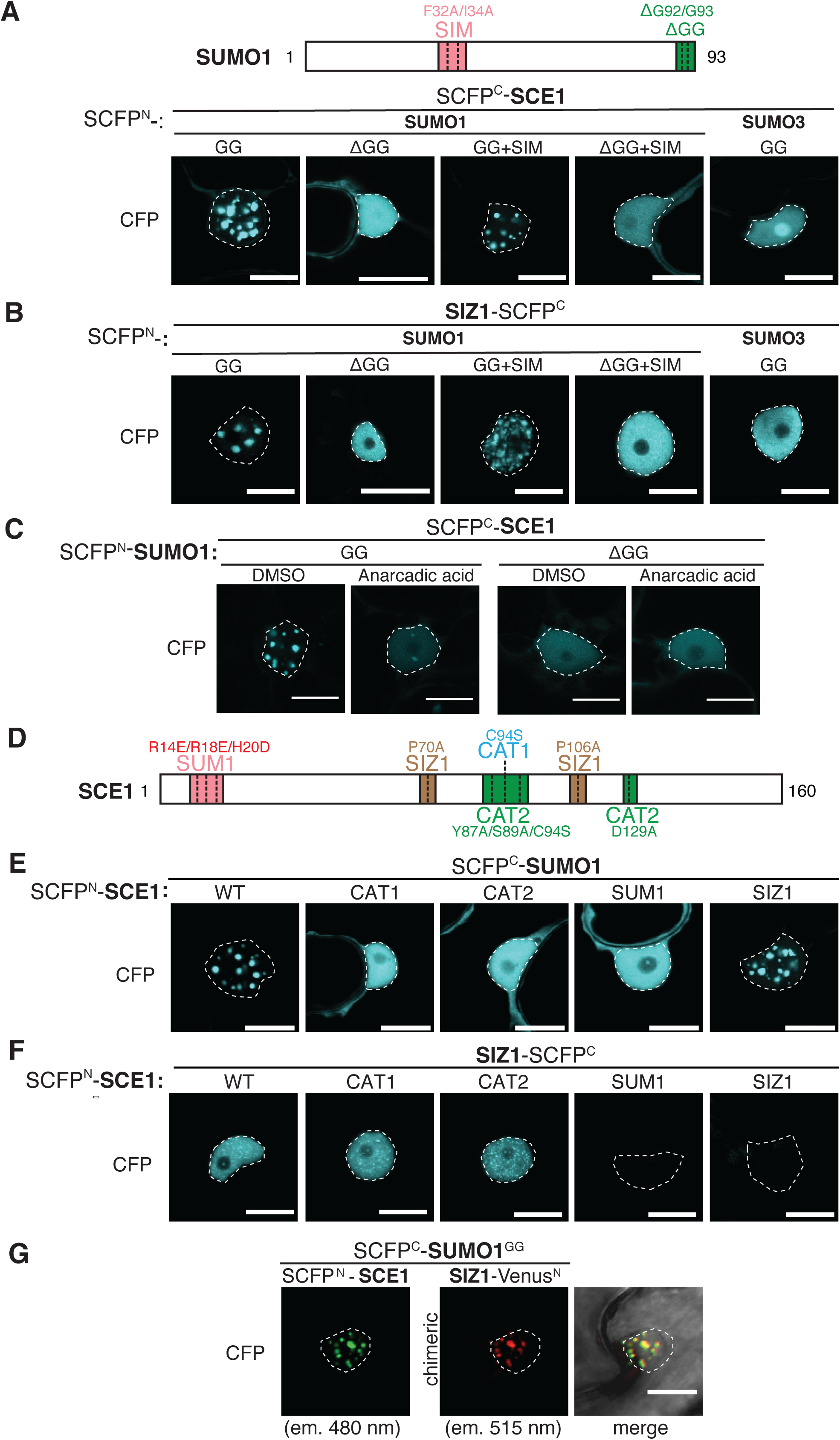
SUMO nuclear body formation depends on SCE1 conjugation activity. A, Nuclear localization pattern of the SUMO1·SCE1 and SUMO3·SCE1 BiFC pairs. Top diagram depicts residues mutated and/or deleted in SUMO1 (GG, mature SUMO; SIM, F32A+I34A; ΔGG, deletion of the C-terminal diGly motif). B, Similar to (A); nuclear localization pattern of the SUMO1/3-SIZ1 BiFC pairs. C, Nuclear localization pattern of the SUMO1·SCE1 couple in response to inhibition of SUMO conjugation using anacardic acid (100 μM in 1% DMSO) 1.5hrs post infiltration. D, Diagram depicting SCE1 mutants and their substitutions. CAT1; catalytic Cys residue mutated; CAT2; binding pocket for the ΨKxE SUMO acceptor motif mutated; SUM1; non-covalent association of SUMO disrupted; SIZ1; SIZ1-binding disrupted. E, Nuclear localization pattern of the SUMO1·SCE1 BiFC pair; SCE1^*CAT1*^, SCE1^*CAT2*^, SCE1^*SUM1*^ do not accumulate in NBs in a BiFC interaction with SUMO1^GG^. F, Nuclear localization pattern of the SCE1·SIZ1 BiFC pair; SCE1^*SUM1*^ and SCE1^*SIZ1*^ fail to interact with SIZ1. G, Multicolour BiFC of SUMO1^GG^, SCE1 and SIZ1 showing complete co-localization in NBs. The micrographs show the nuclear signal of the reconstituted SCFP^N^/SCFP^C^ (SCE1·SUMO1^GG^) and Venus^N^/SCFP^C^ (SIZ·SUMO1^GG^) fluorophores and their merged signals. The two chimeric BiFC couples differ in their excitation and emission spectra. The BiFC pairs in A-F were fused to two halves of SCFP (SCFP^N^+SCFP^C^) with the orientation of the fusions indicated. Scale bar = 20 μm. All micrographs were taken in *N. benthamiana* epidermal leaf cells 2-3 dpi *Agrobacterium* expressing the indicated constructs; nuclei are outlined with white lines. The Supplemental figure S3 depicts for the panels A-F an overlay of the DIC and CFP images of the nuclei shown and a zoom-out (A’-F’) depicting the BiFC signal in the entire cell.

To confirm this notion, we examined if another Arabidopsis SUMO paralogue, SUMO3, could trigger NB assembly in a BiFC interaction with SCE1 or SIZ1. We reasoned that SUMO3 would not trigger NB assembly, as (*i*) it is a poor substrate *in vitro* for SUMO conjugation compared to SUMO1 (Lois et al., 2003), (*ii*) it interacts weakly with SCE1 in the Y2H assay (Supplemental Fig. S1E) and (*iii*) its overexpression in Arabidopsis fails to increase the global level of SUMO1/2 conjugates, while overexpression of SUMO1/2^GG^ or SUMO1/2*^ΔGG^* massively increased these conjugate levels (Van den Burg et al., 2010). Indeed, SUMO3^GG^ failed to activate SCE1 or SIZ1 recruitment to NBs (Fig. 1, A and B and Supplemental Fig. S3), while GFP-tagged SUMO3^GG^ localized to the nucleus and cytoplasm similar to GFP-SUMO1^GG^ (Supplemental Fig. S1, D and F). Additional proof that assembly of these SUMO NBs requires enzymatic activity came from blocking sumoylation using the chemical inhibitor anacardic acid. Anacardic acid directly binds to the E1 enzyme SAE1/2 and prevents formation of the E1∼SUMO intermediate (Fukuda et al., 2009); consequently, it also blocks SUMO transfer onto the E2 protein (E2∼SUMO thioester complex formation). In the presence of anacardic acid, the CFP signal of the SUMO1^GG^·SCE1 pair disappeared from pre-existing NBs within 90 min after adding the inhibitor (Fig. 1C, Supplemental Fig. S3C). Thus, the E1∼ SUMO complex is needed to retain high levels of the SUMO1·SCE1 pair in NBs, meaning that maintenance of these SUMO NBs is a dynamic process.

### Not only SUMO loading (E2∼SUMO) but also the SUMO binding site (E2·SUMO) is critical for SCE1 to assemble in SUMO1·SCE1 NBs

To further discern how these NBs are formed, different Arabidopsis SCE1 variants were used that (i) cannot interact with SUMO or SIZ1, (ii) had lost their catalytic activity (SCE1^*CAT1* and *CAT2*^), or (iii) no longer recognized the ψKxE SUMO acceptor motif in SUMO substrates (SCE1^*CAT2*^) (Mazur et al., 2017) (Fig. 1D; Supplemental Table S1). Studies with the human and yeast SUMO E2 enzymes had identified that the residues Arg14, Arg18 and His20 of SCE1 determine together the binding of SUMO1 to the second non-covalent binding site (the E2**·**SUMO complex) (Bencsath et al., 2002). Likewise, the residues Pro69 and Pro106 of the human Ubc9 (SUMO E2) are essential for the SUMO E2 enzyme to interact with the PIAS family of SUMO E3 ligases (closest homolog of Arabidopsis SIZ1) (Mascle et al., 2013). These five residues are strictly conserved in Arabidopsis SCE1 and other plant homologues of SCE1 (Supplemental Fig. S4), and also for Arabidopsis SCE1 these residues are essential for its interaction with SUMO1 or SIZ1 in the Y2H assay (Mazur et al., 2017). The SCE1^*CAT1*^ mutant still interacted with SUMO1; however, SCE1^*CAT2*^ had lost both its catalytic activity and its non-covalent interaction with SUMO1 (Mazur et al., 2017). See Supplemental Table S1 for an overview of the mutants tested.

Introduction of these mutations in SCE1 did not change its subcellular localization *in planta, i.e.* each variant showed a uniform distribution in both the nucleus and cytoplasm when transiently expressed as GFP-fusion in *N. benthamiana* (Supplemental Fig. S2D). Both SCE1^*CAT1*^ and SCE1^*SUM1*^ appeared to accumulate to higher protein levels *in planta* (Supplemental Fig. S2E), possibly indicating that the stability of wild type SCE1 is negatively impacted by its own enzymatic activity and/or its non-covalent association with SUMO1. Importantly, SCE1^*SIZ1*^ still interacted with SUMO1^GG^ in NBs, but SCE1^*CAT1*^, SCE1^*CAT2*^ and SCE1^*SUM1*^ were not recruited to such NBs while they still interacted with SUMO1^GG^ in the BiFC assay (Fig. 1E and Supplemental Fig. S3E). These findings thus indicate that besides SUMO loading (E2∼ SUMO thioester complex) also the non-covalent SUMO binding (E2·SUMO) is critical for SCE1 to assemble in SUMO1**·**SCE1 NBs.

### The SUMO·SCE1 interface contributes to the SCE1-SIZ1 interaction

In contrast to the SUMO1^GG^·SCE1/SIZ1 couples, we noted that the SCE1·SIZ1 BiFC complex did not aggregate in NBs but rather resided in small nuclear speckles spread across the nucleus (Fig. 1F and Supplemental Fig. S3F). We already showed that the SCE1 residues Pro70 and Pro106 (Supplemental Table S1, SCE1^*SIZ1*^) are essential for SCE1 and SIZ1 to interact in the Y2H assay, while mutating the catalytic site (SCE1^*CAT1*^ and SCE1^*CAT2*^) did not impair their Y2H interaction (Mazur et al., 2017). Also in the BiFC assay, both the SCE1^*SUM1*^ and SCE1^*SIZ1*^ variants failed to interact with SIZ1, while the catalytic-dead variants (SCE1^*CAT1*^ and SCE1^*CAT2*^) still interacted with SIZ1. This suggests that SUMO loading in the catalytic site of SCE1 (SCE1^*CAT1* and *CAT2*^; E2∼ SUMO1) is not required for SCE1 and SIZ1 to interact, while the non-covalent interaction (SCE1^*SUM1*^; E2**·**SUMO1) strengthens this interaction in both the Y2H and BiFC assays (Fig 1F; Mazur et al., 2017). Apparently, the non-covalently bound SUMO acts as a glue in the SCE1-SIZ1 complex or it alters the SCE1 conformation such that it enhances the interaction between SCE1 and SIZ1.

Next, we examined whether the BiFC pairs SUMO1^GG^**·**SCE1 and SUMO1^GG^**·**SIZ1 physically co-localized in one and the same NBs. Thereto, we performed multicolour BiFC (mcBiFC) (Gehl et al., 2009), where SUMO1^GG^ was expressed as fusion protein with SCFP^C^, SCE1 was fused at its N-terminus to SCFP^N^ and SIZ1 was fused at its C-terminus to the Venus residues 1-173 (Venus^N^). Reconstitution of both fluorophores was examined using optical filters that separate CFP (for SCFP^C^-SUMO1^GG^ with SCFP^N^-SCE1) from the chimeric Venus^N^-SCFP^C^ signal (for SCFP^C^-SUMO1^GG^ with Venus^N^-SIZ1). This mcBiFC experiment revealed a complete overlap between the SUMO1^GG^·SCE1 and SUMO1^GG^·SIZ1 signals in NBs (Fig. 1G).

As the SCE1·SIZ1 BiFC pair alone did not form large distinct NBs (Fig. 1F), we tested if the levels of free SUMO1^GG^ could have been limiting, thus preventing the aggregation of the SCE1**·**SIZ1 protein complex into enlarged nuclear structures. We expressed the SCE1·SIZ1 pair together with YFP-tagged SUMO1^GG^ or SUMO1*^ΔGG^* (negative control). The YFP-SUMO1^GG^ signal overlapped with the SCFP signal, but only in a few cells the SCE1**·**SIZ1 BiFC signal shifted from speckles/puncta to enlarged NBs. YFP-SUMO1*^ΔGG^* localized as well to the nucleus, but overall it did not co-localize with the SCE1·SIZ1 BiFC pair in nuclear specks (Supplemental Fig. S5). Thus, the SUMO1-dependent NB enlargement is foremost seen when SUMO is trapped in a BiFC interaction with SCE1 or SIZ1. Possibly, the BiFC system stabilizes a transient protein-protein interaction, which then allows formation of enlarged SUMO NBs. Alternatively, the presence of the intact YFP tag attached to SUMO1^GG^ might cause steric hindrance in the ternary complex.

### SUMO chain formation potentially stimulates formation and enlargement of SUMO NBs

As in mammalian cells SUMO chain formation promotes formation of Promyelocytic Leukemia (PML) NBs (Hattersley et al., 2011), we assessed whether the formation of SUMO NBs depends SUMO chain formation in plants. To block SUMO chain formation, we engineered SUMO1 variants in which Lys9, Lys10 or all seven Lys were replaced by Arg (Supplemental Table S1, SUMO1^*K9R*^, SUMO1^*K10R*^, SUMO1*^K9R+K10R^* and SUMO1*^K7ØR^*). Although Lys10 is the main site for SUMO chain elongation, Lys9 can serve as an alternative site (Colby et al., 2006). Therefore, we also created the SUMO1*^K9R+K10R^* double mutant. To test if these KtoR mutants can still recruit SCE1 to NBs, they were expressed as BiFC pair with SCE1. Two days after *Agrobacterium* infiltration, the NBs formed were reduced in size and number for the KtoR mutants compared to wild type SUMO1, while the diffuse nuclear signal of SCFP had increased (Supplemental Fig. S6A). The SUMO1*^K7Ø^* ·SCE1 pair failed entirely to form NBs two days post infiltration, while SUMO1*^K9R+K10R^*·SCE1 displayed a reduced number of NBs at this time point. These data agree with the proposed role of SUMO chains in NB formation. However, three days post infiltration NBs were found for each KtoR SUMO1 mutant (Supplemental Fig. S6A), albeit the number of cells that contained NBs was less for SUMO1*^K7Ø^*·SCE1 (40-50% for SUMO1*^K7Ø^* compared to 70–100% for the other combinations). Even though, we cannot rule out that these Lys mutations influence protein translation at this time, these data suggest that SUMO chain formation potentially stimulates the targeting of the SUMO1·SCE1 complex to NBs, affecting both their initial formation and subsequent enlargement.

### SUMO1^GG^·SCE1 NBs show complete co-localization with COP1 NBs

Importantly, COP1 co-localizes with SIZ1 in NBs (Kim et al., 2016). Using the Y2H assay, we confirmed that COP1 directly interacts with SIZ1 and not with Arabidopsis SCE1, SUMO1 or SUMO3 (Fig. 2A). We then assessed whether sumoylation of COP1 is needed to co-localize with SUMO1^GG^·SCE1 in NBs. Similar to SIZ1 (Kim et al., 2016), RFP-COP1 co-localized strongly with the SUMO1^GG^·SCE1 BiFC pair in NBs (Fig. 2, B and C and Supplemental Fig. S6C). Importantly, both the SUMO1*^ΔGG^*·SCE1 and SUMO1^GG^**·**SCE1^*CAT1*^ pairs (Fig. 2, B and C) still failed to localize to NBs regardless of RFP-COP1 overexpression. Thus, co-localization of SCE1 and COP1 in NBs requires again SCE1 catalytic activity. Co-expression of RFP-COP1 with different SUMO1^*KtoR*^·SCE1 BiFC pairs revealed that all these SUMO1^*KtoR*^·SCE1 variants strongly co-localized with COP1 in NBs irrespective of the lysine mutations (Supplemental Fig. S6B). Next, we tested if the COP1 SUMO acceptor site (Lys193) is important for this co-localization. Introduction of the K193R mutation (COP1^*SUMO*^) reduced the overlap between the COP1 and SUMO1·SCE1 NBs (Fig. 3, B and C, Supplemental Fig. S7A), but it did not suppress the COP1-SIZ1 protein-protein interaction in the Y2H assay (Fig. 3, A and D) suggesting that residues other than Lys193 promote the interaction between the E3 ligases SIZ1 and COP1.

**Figure 2.**
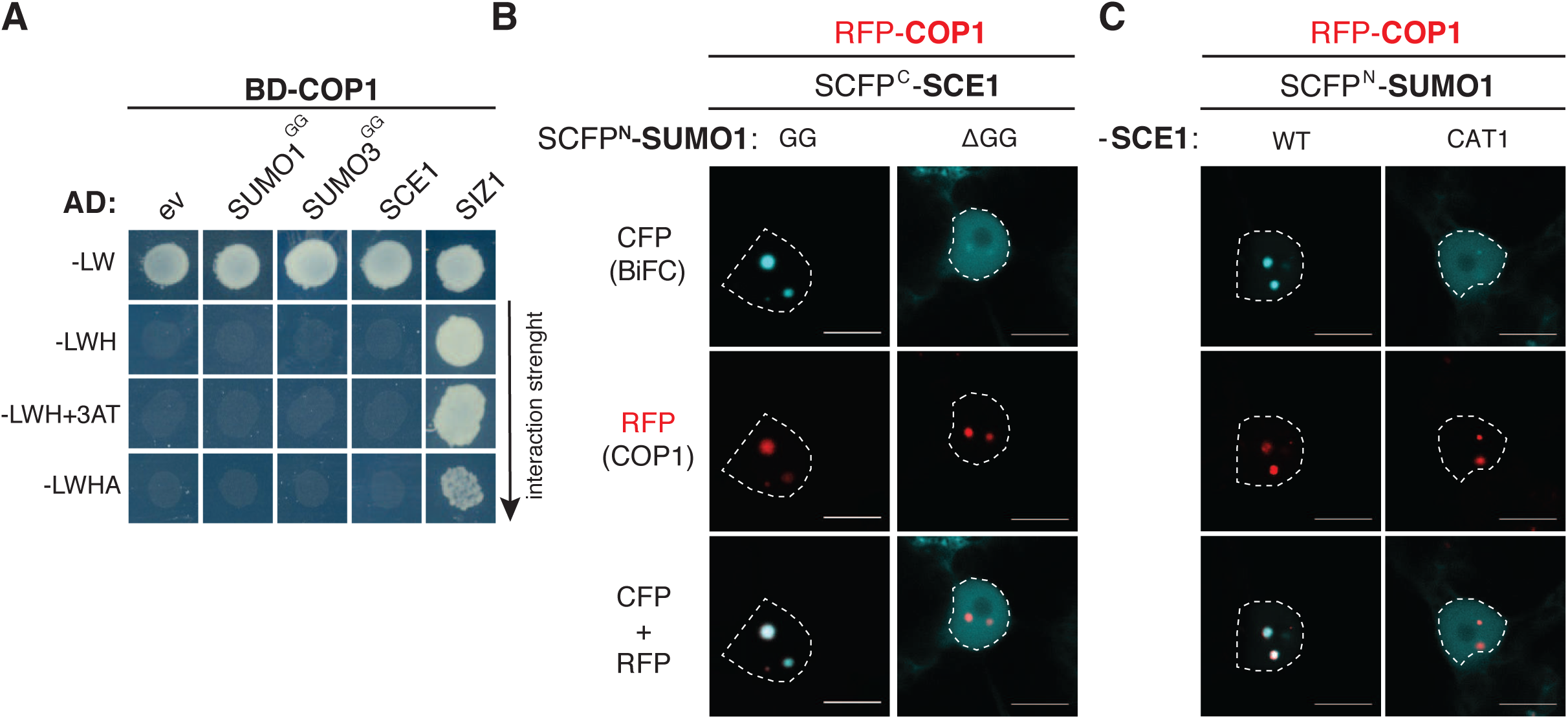
The SUMO1·SCE1 BiFC pair co-localizes with COP1 in nuclear bodies when catalytically-active. A, Yeast two-hybrid assay expressing COP1 as GAL4 BD fusion protein and the SUMO (machinery) proteins were fused to the GAL4 AD-domain. Yeast growth was scored 3 days after incubation on selective media at 30°C (-LW, -LWH, -LWH+ 1mM 3AT, -LWHA). B and C, Nuclear localization pattern of the SUMO1·SCE1 BiFC pair in cells overexpressing RFP-COP1. Only wild type SCE1 (WT) with SUMO1^GG^ (GG) co-localizes as BiFC couple with COP1 in NBs. B: ΔGG, conjugation-deficient SUMO variant; C: CAT1, Catalytically-dead SCE1. Micrographs show from top-to-bottom the reconstituted BiFC signal, RFP-COP1 and their merged signals. Conditions were identical to Fig 1. Scale bar = 10 μm.

**Figure 3.**
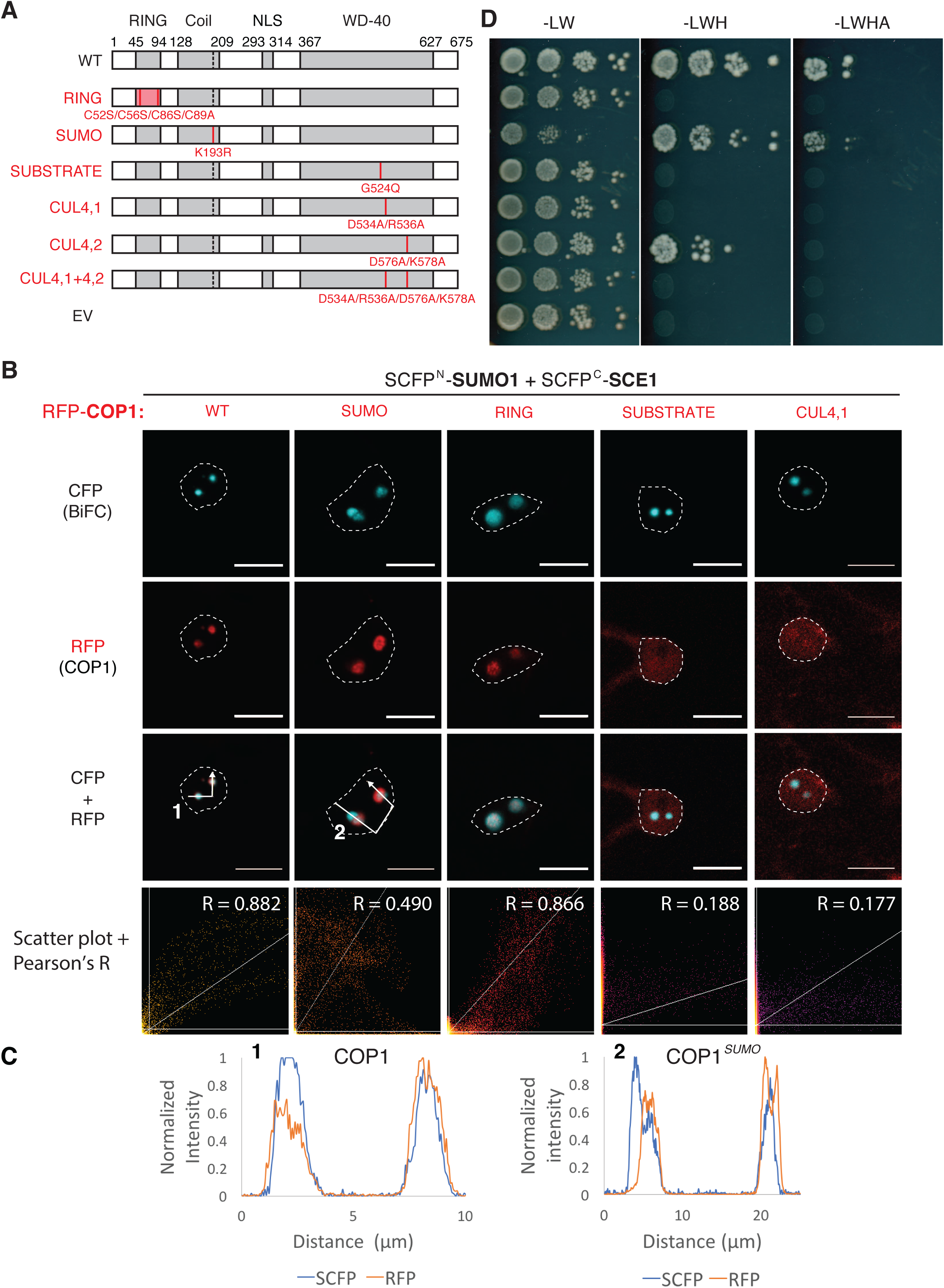
The COP1-SIZ1 interaction in NBs depends on both the substrate binding pocket and the SUMO acceptor site in COP1. A, Diagram depicting COP1 variants and the protein-protein interaction domain disrupted (left side, red text) with their mutations shown: *RING,* RING Zn2+-finger binding domain mutant; *SUMO*, loss of SUMO acceptor site; *SUBSTRATE*, mutation in the substrate binding groove; *CUL4*, mutations in the CUL4-binding ‘WDRX’ motifs. B, Nuclear localization pattern of the SUMO1·SCE1 BiFC couple in cells overexpressing functional mutants of RFP-COP1. Micrographs depict from top-to-bottom the BiFC CFP signal, RFP-COP1, and their combined signals. Bottom row depicts a scatter plot of the RFP/CFP signal intensities per pixel and their Pearson’s correlation coefficient (R). In particular, the COP1 SUMO acceptor site (COP1^*SUMO*^) promotes colocalization of SCE1·SUMO1 and COP1 in the same NBs. Conditions were identical to Fig 1. Scale bar = 10 μm. C, Normalized intensity profiles depict the fluorescence signal intensities for SCFP: (BiFC pair) and RFP-COP1: co-expression of (1) WT COP1 or (2) COP1^SUMO^; the profiles follow the white arrows depicted in panel B. Note the profiles of RFP-COP1^*SUMO*^ and SUMO·SCE1 CFP signals poorly overlap in (2). D, Mapping of the SIZ1-interaction site in COP1 using the yeast 2-hybrid assay. The COP1 variants were fused to the GAL4 BD; SIZ1 was fused to the GAL4 AD-fusion. Yeast growth was scored 3 days after incubation on selective medium at 30°C.

Structural studies revealed that substrates of COP1 interact with it via VP peptide motifs that dock in the central groove of the COP1 WD-40 propeller head (Uljon et al., 2016). The mutation Q529E (the causal mutation in the *cop1-9* allele, hereafter COP1^*SUBSTRATE*^) is positioned in this VP-binding groove and was shown to disrupt the binding of different COP1 substrates (Holm et al., 2002). Importantly, this COP1^*SUBSTRATE*^ variant fails to form NBs when expressed as a GFP-fusion protein (Stacey et al., 1999). COP1 also interacts with the DBB1-CUL4 E3 ubiquitin ligase complex via two WDRX motifs; these WDRX motifs are also located in this WD-40 domain near the VP-binding groove (Chen et al., 2010).

We used these COP1 variants to test whether the capacity of COP1 to bind substrates or CUL4 (i) determines its targeting to NBs and (ii) related its recruitment to SUMO NBs. First, we tested if these COP1 variants still interacted with SIZ1 in the Y2H assay (Fig. 3D). Except for COP1*^SUMO^,* all the other mutants failed to interact with SIZ1 (Fig. 3D). When fused to RFP, COP1^*SUMO*^ and COP1^*RING*^ still formed NBs in *N. benthamiana*, while the variants COP1^*SUBSTRATE*^, COP1*^CUL4,1^* and COP1*^CUL4,1 and 4,2^* failed to localize to NBs *in planta* (Fig. 3C, Supplemental Fig. S7B). Mutations in the WD-40 domain thus suppressed formation of COP1 NBs. To quantify the degree of co-localisation between the COP1 variants and the SUMO1^GG^·SCE1 BiFC signal, the RFP/CFP pixel intensities were depicted in a scatter plot. This yielded a strong positive correlation between the signal intensities of wild type RFP-COP1 and the SUMO1^GG^·SCE1 CFP signal (Pearson’s R = 0.882). The localization of RFP-COP1^*RING*^ also correlated strongly with the SUMO1^GG^·SCE1 CFP signal (Pearson’s R = 0.866). The degree of co-localization was less for RFP-COP1^*SUMO*^ (Pearson’s R = 0.490), while localization of the variants RFP-COP1^*SUBSTRATE*^, -COP1*^CUL4,1^* and -COP1*^CUL4,1 and 2^* did not correlate with the SUMO1^GG^·SCE1 signal (Fig. 3B, and Supplemental Fig. S7B; Pearson’s R = 0.188, 0.177 and 0.148, respectively).

RFP-COP1 also co-localized with the SUMO1·SIZ1 pair in NBs (Fig. 4A). Their co-localization in NBs required an intact substrate-binding pocket in COP1 (COP1^*SUBSTRATE*^). Similar to the SUMO1·SCE1 BiFC pair, the SUMO acceptor site in COP1 (COP1^*SUMO*^) contributed to the overlap between the SUMO1·SIZ1 and RFP-COP1 signals in NBs, while disruption of COP1 ubiquitin ligase activity (COP1^*RING*^) had no apparent effect on targeting of the SUMO1·SIZ1 complex to these COP1 NBs (Fig. 4A). Thus, the substrate binding pocket of COP1 appears to be the main determinant for both COP1 recruitment to NBs *and* its binding with SIZ1.

**Figure 4.**
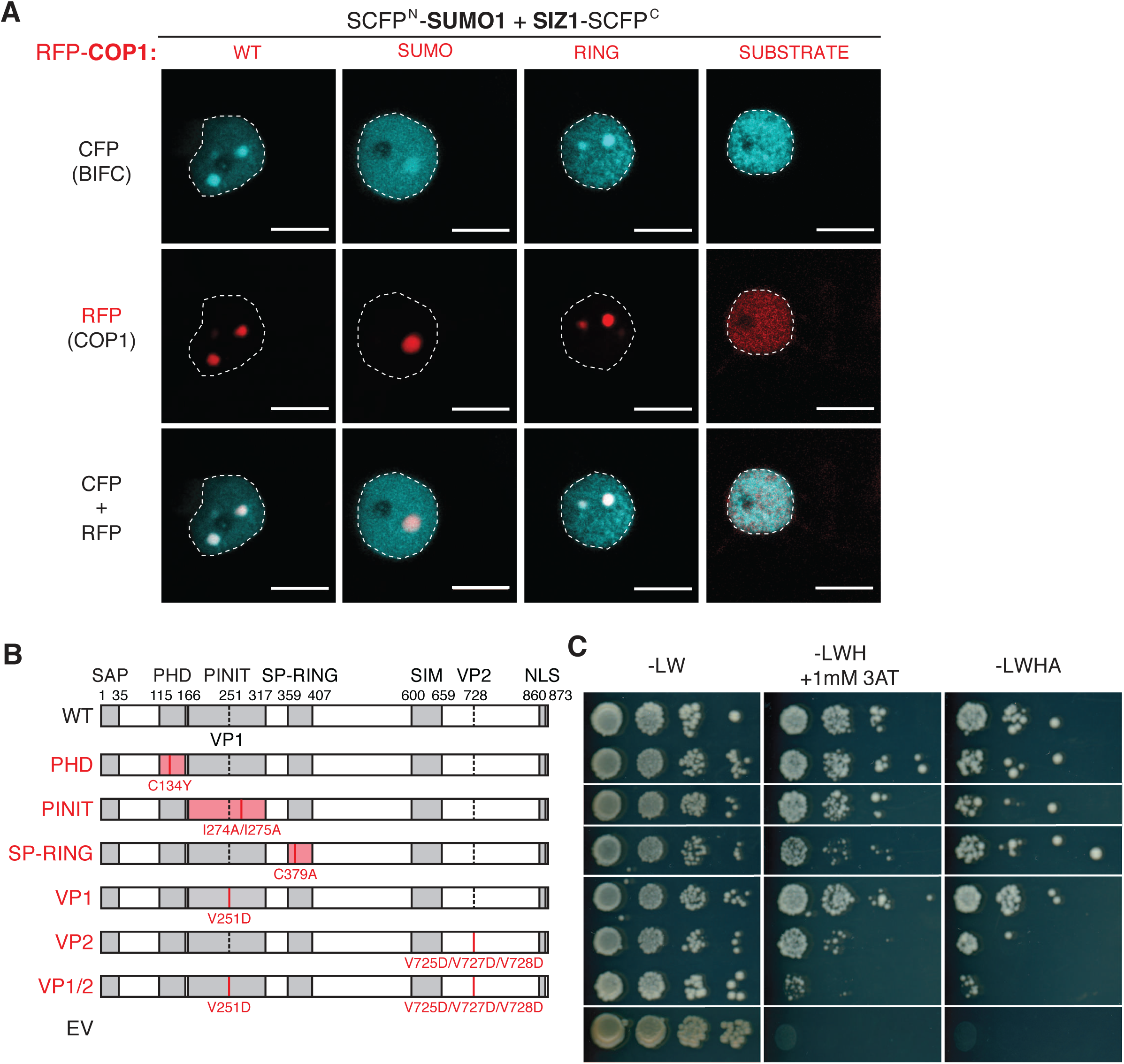
Formation of the multiparty COP1+SUMO1·SIZ1 NBs depends on the COP1 substrate binding pocket that apparently recruits SIZ1 via VP-motifs. A, Nuclear localization pattern of the SUMO1·SIZ1 BiFC couple in cells overexpressing RFP-COP1 variants. The COP1 substrate-binding pocket is essential for COP1 NB formation, while co-localization of the SUMO1^GG^·SIZ1 BiFC pair with RFP-COP1 depends on both the SUMO acceptor site (SUMO) and the substrate binding pocket of COP1 (SUBSTRATE). See Fig. 4A for details on the COP1 variants. Micrographs show from top-to-bottom the BiFC signal, RFP-COP1, and their merged signals. Conditions were identical to Fig 1. Scale bar = 10 μm. B, Diagram depicting SIZ1 variants and the protein-protein interaction domain disrupted (left side) by the mutations introduced: PHD and PINIT, both reduced substrate binding; SP-RING, no interaction with SCE1; VP1 and VP2, putative interacting motifs for the COP1 substrate binding groove. C, Mapping of the COP1-interaction site in SIZ1 using the yeast 2-hybrid assay. The COP1-SIZ1 interaction depends on an intact SP-RING and VP2 domain in SIZ1. Similar to Fig. 3, SIZ1 variants were fused to GAL4 AD-fusion, while COP1 was fused to GAL4 BD; Yeast growth was scored after 3 days at 30°C.

### SIZ1 contains two VP domains important for COP1 binding

To map the COP1-binding site in SIZ1, a series of SIZ1 mutants was prepared. In yeast both the PHD and PINIT domain are essential for *Sc*SIZ1-directed sumoylation of the substrate PCNA (Proliferating cell nuclear antigen) (Yunus and Lima, 2009). Loss-of-function mutations in these two domains in Arabidopsis SIZ1 did not compromise the interaction with COP1 in the Y2H assay. Disruption of the SP-RING domain, which is needed to recruit the E2∼SUMO thioester into a complex with its substrate PCNA (Yunus and Lima, 2009), impaired the interaction between SIZ1 and COP1 (Fig. 4, B and C). SIZ1 also contains two VP motifs (VP1:Val251 and VP2:Val725/728), which could explain why SIZ1 is recognized by COP1 as a substrate. Both motifs were mutated by replacing Val for Asp residues. Mutating VP1 did not suppress the SIZ1-COP1 interaction, while mutating VP2 reduced this interaction. Furthermore, mutating both VP motifs disrupted the interaction entirely (Fig. 4, B and C). Thus, the SP-RING domain of SIZ1 contributes to the interaction with COP1, but the VP motifs combined are essential for this interaction.

To determine if SCE1 and SIZ1 can already reside in COP1 NBs independent of their SUMO-BiFC interaction, we co-expressed GFP-tagged SCE1 or SIZ1 together with RFP-tagged COP1 or COP1^*SUMO*^. For both proteins (SCE1 and SIZ1), the GFP signal was enriched in RFP-COP1 NBs (Fig. 5A). Moreover, their targeting to COP1 NBs was significantly less when they were co-expressed with RFP-COP1^*SUMO*^ (Figs. 5, A and B) and these latter GFP-SIZ1/RFP-COP1^*SUMO*^ NBs also had an amorphous shape (asterisk in Fig. 5A). Combined, these data suggest that the SUMO acceptor site in COP1 contributes to the recruitment of SCE1 and SIZ1 to COP1 NBs.

**Figure 5.**
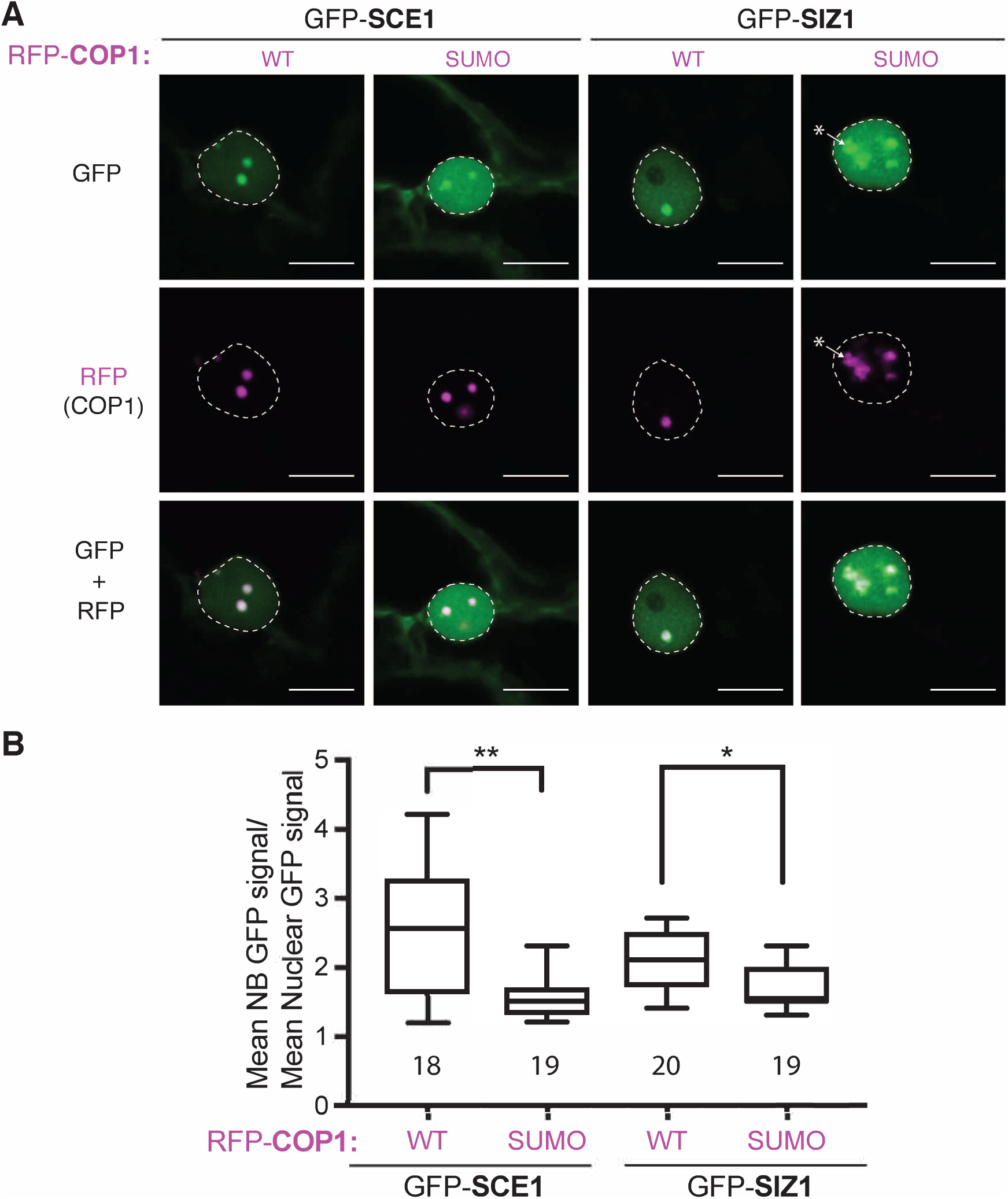
Co-localization of GFP-tagged SCE1/SIZ1 with RFP-COP1 NBs is compromised when the SUMO acceptor site is mutated in COP1. A, Nuclear localization pattern of GFP-tagged SCE1 and SIZ1 is modulated by loss of the SUMO acceptor site (Lys193Arg, COP1^*SUMO*^) in RFP-COP1. Remarkably, GFP-SIZ1 forms together with RFP-COP1^*SUMO*^ amorphous NBs (marked with *). Micrographs from top-to-bottom: GFP, RFP and their merged signals. Scale bar = 10 μm. B, Quantification of the average GFP signal intensity in the NBs per nucleus divided by the average fluorescence signal in the nucleus (with the data of 3 biological replicates pooled with at least 5 nuclei per replica). Total number of nuclei analyzed is shown. Significant differences were detected using an unpaired Student’s T-test assuming unequal variances; ** p < 0.01, * p < 0.05. Conditions were identical to Fig. 1.

### Recruitment of SCE1 to COP1 NBs requires the *SIZ1* gene in Arabidopsis

As the native proteins are still present in the *N. benthamiana* BiFC experiments, we shifted to particle bombardment in Arabidopsis leaf epidermal cells. Using the Arabidopsis mutants *cop1-4, siz1-2,* and *sumo1*;*amiR-SUMO2*, we genetically inferred whether the endogenous proteins are required for (co-)localization of SCE1 and COP1 in NBs. The different constructs where shifted to high expression vectors suitable for particle bombardment (Walter et al., 2004) in which SCE1 and SUMO1^GG^/SUMO1*^ΔGG^* were tagged at their N-terminus with the BiFC halves ^N^YFP and ^C^YFP, respectively. Similar to the *N. benthamiana* data, the SUMO1^GG^·SCE1 combination formed NBs in Arabidopsis, while SUMO1*^ΔGG^*·SCE1 interacted but this combination displayed a uniform BiFC signal across the nucleus (Fig. 6A). Targeting of SUMO1^GG^·SCE1 to NBs did not change when we the BiFC protein complex was expressed in the genotypes *cop1-4* or *siz1-2*, confirming that SUMO conjugation is the main force behind SUMO NB formation. To assess genetically if sumoylation activity is essential for COP1 NB formation, GFP-COP1 was expressed in wild type Arabidopsis (Col-0), *siz1-2* and *sumo1*;*amiR-SUMO2* using particle bombardment. Irrespective of these three genetic backgrounds, GFP-tagged COP1 localized to NBs in the Arabidopsis plants (Fig. 6B). This corroborates our notion that SUMO conjugation activity *per se* is not the main driving force for COP1 aggregation in NBs. To demonstrate that the co-localization of SCE1 and COP1 in NBs depends on SIZ1, RFP-SCE1 and GFP-COP1 were transiently co-expressed in wild type Arabidopsis and *siz1-2* using particle bombardment. As expected, in wild type bombarded Arabidopsis rosettes RFP-SCE1 was enriched in GFP-COP1 NBs, while in the *siz1-2* background RFP-SCE1 was absent from these COP1 NBs (Fig. 6, B and C). Thus, recruitment of SCE1 and COP1 to one and the same NB requires the endogenous SIZ1 protein to be present.

**Figure 6.**
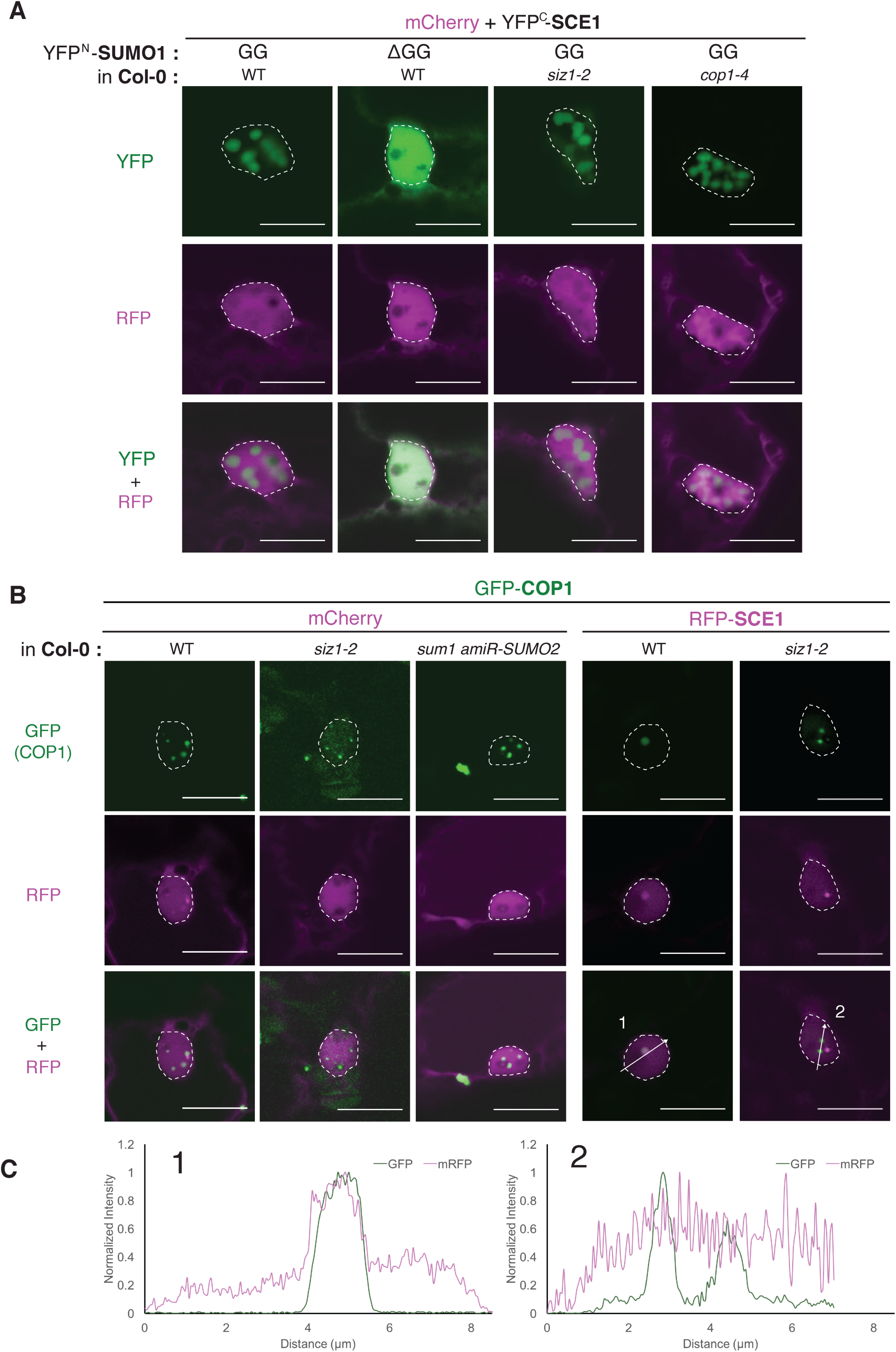
SUMO conjugation triggers also NB formation in Arabidopsis, which depends genetically on *SIZ1* for recruitment to COP1 NBs. A, Nuclear localization pattern of SCE1·SCE1 BiFC pair in Arabidopsis (wild type, *siz1-2* or *cop1-4*) using particle bombardment. The BiFC constructs and the used Arabidopsis genotypes are indicated at the top; to detect the transformed cells, mCherry was co-bombarded. Again SUMO1^GG^·SCE1 is effectively targeted to NBs in Arabidopsis independent of the *SIZ1* and *COP1* genes. The micrographs show from top to bottom the YFP, RFP and the merged signal; nuclei are outline with a white line. 4-week-old Arabidopsis rosettes were bombarded. Two days post bombardment, plant material was transferred to a dark box and 1 day later the fluorescence signals were examined. B, Nuclear localization pattern of GFP-COP1 and RFP-SCE1 in Arabidopsis (wild type, *siz1-2* or *sumo1;amiR-SUMO2*) using particle bombardment. To detect the transformed cells, either mCherry was co-bombarded or RFP-SCE1 was used. COP1 localizes to NBs independent of SIZ1 or SUMO conjugation. Micrographs with nuclei with GFP-COP1 containing NBs are shown. Scale bar = 10 μm.

### Simultaneous recruitment of CRY1/2 and SUMO1·SCE1 to COP1 NBs

CRY1 and -2 are blue light receptors that inhibit COP1 activity indirectly via the SPA proteins or directly, respectively, after blue light exposure (Wang et al., 2001; Liu et al., 2011; Holtkotte et al., 2017). Fluorescent protein fusions of CRY2 localize to NBs after blue light exposure (Mas et al., 2000). As, at least to our knowledge, neither CRY1 nor CRY2 are sumoylated, their subcellular localization might correlate with COP1, but not with the SUMO conjugation enzymes. CRY1-GFP and CRY2-GFP were co-expressed with RFP-COP1, RFP-SCE1 and SIZ1-RFP in *N. benthamiana* cells. CRY2-GFP localized in our system in NBs and CRY1-GFP localized to the whole nucleus. As expected, both CRY1 and CRY2 were recruited to mRFP-COP1 NBs (Supplemental Fig. 8, A and B). SIZ1-mRFP or mRFP-SCE1 co-expressed with CRY1/CRY2-GFP were evenly distributed in the nucleus and did neither alter CRY1-nor CRY2-GFP localization. Importantly, SCE1 and SIZ1 were not recruited to CRY2-GFP NBs. Next, the SUMO1^GG^·SCE1 BiFC pair was co-expressed with CRY1-mRFP or CRY2-mRFP. Neither CRY1-mRFP nor CRY2-mRFP was recruited to the SUMO1^GG^·SCE1 NBs (Fig. 7). However, when YFP-COP1 was co-expressed in the same cells, all components co-localized to the same NBs. Thus, CRY2 NBs and SUMO1^GG^·SCE1 NBs are different entities that coincide in COP1 NBs.

**Figure 7.**
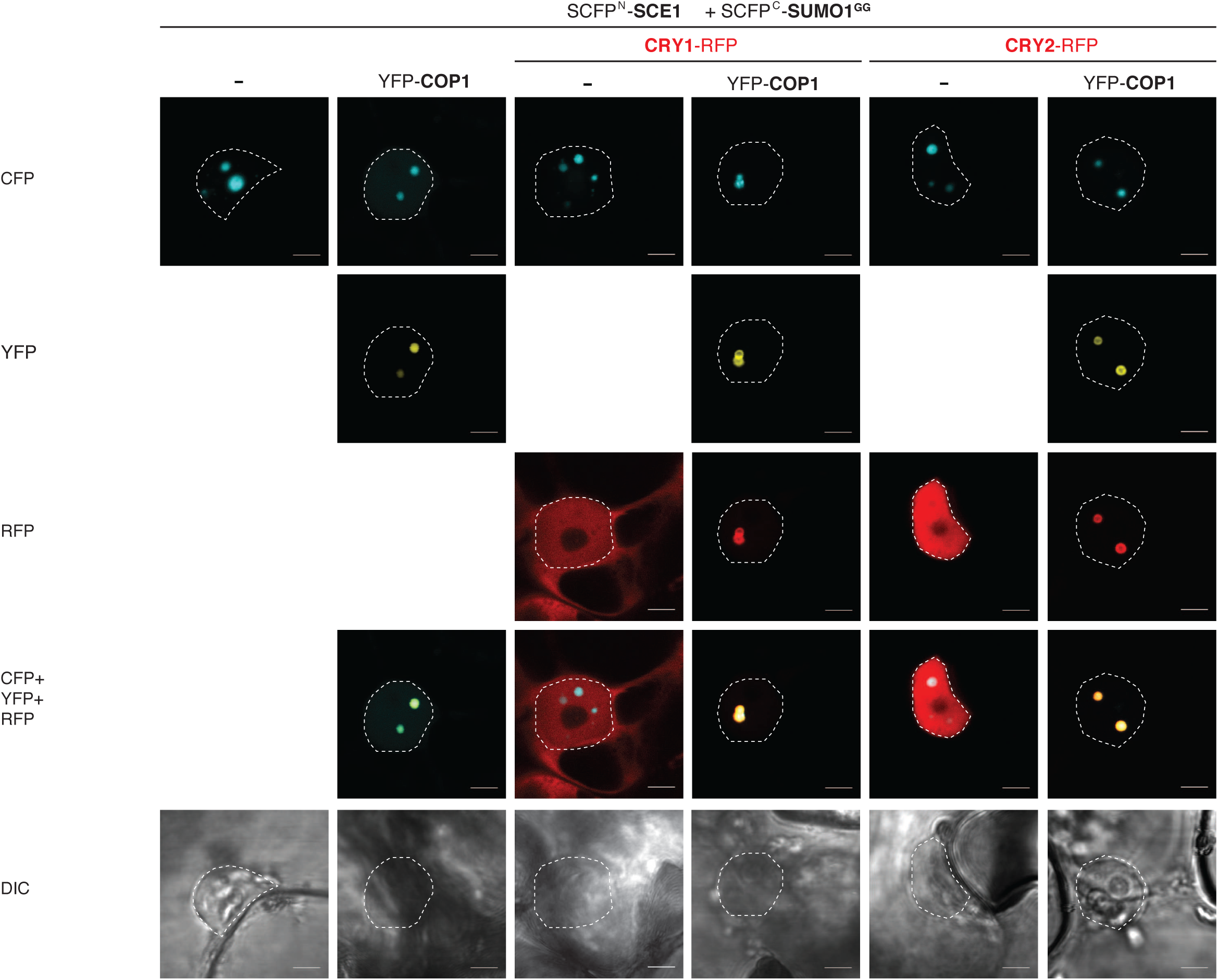
CRY1-mRFP or CRY2-mRFP and the SUMO1·SCE1 BiFC complex, meet in COP1 NBs. Micrographs of cells transiently expressing the SCFP^N^-SCE1 and SCFP^C^-SUMO1^GG^ with or without YFP-COP1 and either CRY1-mRFP or CRY2-mRFP 3 days after Agrobacterium infiltration. Note that neither CRY1-mRFP nor CRY2-mRFP are recruited to SCE1·SUMO1^GG^ NBs. Note that SCE1·SUMO1^GG^ and CRY1-mRFP or CRY2-mRFP are recruited together to YFP-COP1 NBs. Scale bar = 10 μm

## Discussion

Here we characterized the assembly of the ternary complex between SUMO1, SCE1 and SIZ1 in NBs. SCE1 activity (i.e. SCE1∼SUMO thioester) and the non-covalent association of SUMO with SCE1 are both required to target this complex to NBs (Fig. 1, A and E). Strikingly, COP1 fully co-localizes with these SUMO NBs. COP1 is a master regulator of skoto-and thermomorphogenesis that translocates from the cytosol to the nucleus at night-time and at high temperature (28°C) conditions (Höcker, 2017; Park et al., 2017). Once nuclear localized, COP1 targets a set of substrates, mainly transcription factors (TFs) that act as positive regulators of light signaling, for degradation. We found that mutating the COP1 SUMO acceptor lysine (Lys193) reduced the overlap between COP1 and the BiFC pair SCE1·SUMO1 in NBs albeit that both still predominantly localized to NBs (Fig. 3, B and C, Supplemental Fig. S7A). In support, GFP-tagged SIZ1 and SCE1 proteins showed reduced co-localization with COP1 NBs when co-expressed with a COP1 variant that lacks this acceptor lysine (K193R, COP1^*SUMO*^) compared to wild type COP1 (Fig. 5). Moreover, COP1 interacted physically with SIZ1 and not with SCE1, SUMO1 or SUMO3 in the Y2H assay. Importantly, COP1 is a SIZ1-dependent sumoylation substrate and its sumoylation enhances its biochemical activity, resulting in ubiquitylation and degradation of SIZ1 and other substrates, which in turn results in enhanced plant growth (Kim et al., 2016; Lin et al., 2016; Kim et al., 2017). Structural studies had exposed that COP1 substrates dock in a central groove of WD-40 propeller head of COP1 via short VP peptide motifs (Uljon et al., 2016). We identified two of such VP motifs in SIZ1 that together control SIZ1 binding to COP1 (Fig. 4, B and C). Genetically, targeting of SCE1 to COP1 NBs also requires a functional *SIZ1* gene in Arabidopsis (Fig. 6, B and C). Combined, these data signify that SIZ1 is recognized by COP1 as a *bona fide* substrate, likely via its two VP motifs, and that in turn SIZ1 can recruit SCE1 to COP1 NBs.

### Substrate binding drives COP1 to NBs independent of SIZ1

Structural studies had exposed that the C-terminal half of COP1, including its WD-40 domain, is important for substrate binding, but also for its localization to NBs (Stacey et al., 1999; Stacey and von Arnim, 1999; Holm et al., 2002; Uljon et al., 2016). In particular, the amino acid change G524Q found in the *cop1-9* allele is located at the central substrate-binding groove of the WD-40 propeller head and this mutation prevents COP1 targeting to NBs (Stacey et al., 1999; Uljon et al., 2016, Fig. 3B). In contrast, sumoylation of COP1 at Lys193 and the ubiquitin ligase activity of its RING domain (Stacey et al., 1999; Stacey and von Arnim, 1999) are not critical for COP1 aggregation in NBs (Fig. 3B). Using particle-bombardment in Arabidopsis mutants, we established genetically that COP1 resides in NBs independent of SIZ1 (Fig. 6B). As well, the BiFC pair SCE1·SUMO1^GG^ still aggregated in NBs when transiently expressed in *siz1-2* or *cop1-4* (a strong *COP1* allele with little residual COP1 activity remaining). Thus, recruitment of COP1 or SUMO·SCE1 to NBs does not depend on the physical interaction between COP1 and SIZ1, while their collision in the same NBs requires SIZ1 as bridge protein.

### Sumoylation and blue light signalling coincide in COP1 bodies

The blue light receptors CRY1 and CRY2 inhibit COP1 activity in a blue-light dependent manner (Wang et al., 2001). The interaction of CRY2 and COP1 is direct, while CRY1 uses SPA1 as a bridge protein (Wang et al., 2001; Liu et al., 2011; Holtkotte et al., 2017). Directly after blue light exposure, CRY2 is rapidly degraded. COP1 is only partially responsible for this light-dependent degradation of CRY2 (Shalitin et al., 2002). We observed that in the absence of YFP-COP1 overexpression neither CRY1 nor CRY2 co-localize with SCE1, SIZ1 or the SUMO-SCE1 BiFC complex (Fig. 7 and Supplemental Fig. S8). However, when COP1 is present all these proteins are targeted to the same NBs. This suggests that the first steps in the blue light response are sumoylation independent, thereafter all components are apparently dragged to the same NBs for downstream responses. As CRY1 is not and CRY2 is for its degradation only partially dependent on COP1, their action might, however, affect or be affected by COP1 sumoylation. As sumoylation enhances COP1 activity (Lin et al., 2016), CRY1 or CRY2 might either prevent COP1 sumoylation or make COP1 insensitive to CRY1 or CRY2 interference.

### Enzymatic activity of SCE1 drives localization of SUMO1·SCE1 to NBs

Our data on the molecular interactions between SUMO1, SCE1, and SIZ1 support that they adopt a similar ternary complex in Arabidopsis, as previously reported for their yeast and human counterparts (Bencsath et al., 2002; Bernier-Villamor et al., 2002; Reverter and Lima, 2005; Mascle et al., 2013; Sekhri et al., 2015; Mazur et al., 2017). Recruitment of this ternary complex to NBs depends tightly on their intermolecular interactions in *N. benthamiana* and *Arabidopsis*. Furthermore, we established that the SIM binding cleft around Phe32 (Colby et al., 2006) is functionally conserved in Arabidopsis SUMO1 and that it enforces the interaction of SUMO with SCE1/SIZ1 (Fig. 1A). Mutations in the SIM did not suppress assembly of the SUMO·SCE1 and SUMO·SIZ1 NBs, while conjugation-deficient variants of SUMO1 failed to aggregate in these SUMO NBs (Fig. 1, A and B). Thus, a key factor for SUMO NB assembly is conjugation activity. Two lines of evidence argue that the SUMO-SCE1/SIZ1 NBs are not an artefact of the BIFC system or caused by unspecific aggregation of (misfolded) proteins, but rather require active formation and/or maintenance. First, pre-existing NBs disappear within 90 minutes after inhibition of the SUMO E1 enzyme by anacardic acid. Second, NB assembly requires the catalytic site of SCE1 to be intact.

Our data suggest that SUMO chain formation is not essential but might promote formation of the SUMO1·SCE1 NBs (Supplemental Fig. S6A). The non-covalent association of SUMO with SCE1 was reported to be important for SUMO chain formation (Knipscheer et al., 2007). Our data support a mechanistic model in which thioester (E2∼ SUMO) formation is essential, while the non-covalent-association of SUMO (E2**·**SUMO) has a dual role in the redistribution of Arabidopsis SUMO1 and SCE1 to NBs. This is reminiscent to the mechanism by which SUMO drives NB formation in yeast and human cells, e.g. Pc2 (Polychrome 2) in Polycomb group bodies and PML bodies (Yang and Sharrocks, 2010; Jentsch and Psakhye, 2013). Not only the SUMO conjugation machinery localizes to NBs (this study), but also a substantial number of Arabidopsis SUMO substrates and several SUMO proteases have been reported to localize to nuclear speckles/bodies/puncta. For example, the SUMO protease OTS2 (OVERLY TOLERANT TO SALT2) and the putative SUMO-targeted Ubiquitin E3 ligases STUbL1 and STUbL4 were reported to localize to undefined nuclear puncta, resembling the NBs here seen (Conti et al., 2008; Elrouby et al., 2013). Future studies should address if all these bodies reflect one and the same nuclear structure.

### Coincidental or functional: the overlap between COP1 and SIZ1 substrates?

This study strengthens the notion that COP1 and sumoylation go *hand-in-hand* in controlling each other’s activities while simultaneously targeting shared substrates. Interestingly, many of the COP1 substrates and interactors are also (putative) SUMO conjugation targets (phyB, DELLAs, HFR1 (LONG HYPOCOTYL IN FAR RED 1), LAF1 (LONG AFTER FAR-RED LIGHT 1), HY5 (ELONGATED HYPOCOTYL 5), and HYL (HY5 HOMOLOGUE)) (Ballesteros et al., 2001; Seo et al., 2003; Conti et al., 2014; Sadanandom et al., 2015; Tan et al., 2015; Mazur et al., 2017). Jointly, these COP1 substrates control at the transcriptional and post-translational level the activity of the PIFs, which in turn control the light and temperature-sensitive growth pathways (Paik et al., 2017). Moreover, the TF ABI5 (ABA-INSENSITIVE 5), a known interactor of SIZ1 and a SUMO substrate, was also found to translocate to COP1-containing NBs in the presence of the regulator AFP (ABI FIVE INTERACTING PROTEIN) (Lopez-Molina et al., 2003; Miura et al., 2009). Disruption of a putative SUMO acceptor site in the TF LAF1 prevented its translocation to nuclear speckles (Ballesteros et al., 2001), while the TF HFR1 was shown to interact with the bacterial SUMO protease XopD (*Xanthomonas* outer protein D from the bacterium *Xanthomonas*) in nuclear speckles (Tan et al., 2015). Translocation of HFR1 and ABI5 to NBs was in each case associated with their degradation, which provides again a link between COP1 and sumoylation in the regulation of nuclear processes. Considering that SIZ1 controls both skoto- and thermomorphogenesis (Lin et al., 2016; Hammoudi et al., 2018), the recruitment of SIZ1 to COP1 bodies might suggest the existence of a (transient) SUMO conjugation wave in these bodies with unknown physiological and biochemical consequences in a normal diurnal dark/light cycle. As previously shown, SIZ1 is needed for hypocotyl elongation in both darkness and/or high temperature conditions and the *siz1-2* mutation delays and weakens significantly the transcription response of substantial proportion of the PIF4 genomic targets (Lin et al., 2016; Hammoudi et al., 2018). Thus, in addition to having a role in plant immunity, SIZ1-dependent sumoylation rapidly sees the light as an important positive regulator of dark/high temperature-induced growth responses.

## Materials and methods

### Construction mutants and vectors for yeast two-hybrid interactions

All molecular techniques were performed using standard methods (Sambrook and Russel, 2001). Primers (synthesized by Eurofins genomics) are listed in Supplemental Table S2. Information on clones with primers can be found in Supplemental Table S3. Primers containing the *attB1* and *attB2* recombination sites were used to amplify the CDS sequences (see Supplemental Table S2). The PCR products were cloned into the pDONR207 or pDONR221 using BP Clonase II (Thermo Fischer) and checked by sequencing. The inserts were transferred to destination vectors using Gateway LR Clonase II (Thermo Fischer) and the clones were re-sequenced. For the GAL4 BD/AD-fusion yeast-two-hybrid constructs, the cDNA clones were introduced in pDEST22/pDEST32 (Thermo Fischer). As the interactions of SUMO1·SCE1 were weak in the pDEST system (due to low expression levels, Supplemental Fig. S1B), these proteins were expressed with pGBKT7/pGADT7 (Clontech). CRY1 (G12079) and CRY2 (G19559) cDNA clones were obtained from the Arabidopsis Biological Research Centre. For *in planta* protein localization, the CDS were introduced in the destination plasmids pGWB452 (N-terminal GFP tagging), pGWB442 (N-terminal YFP tagging) or pGWB655 (N-terminal mRFP-tagging) (Nakagawa et al., 2007). For BiFC studies, we used the destination vectors pSCYNE(R) (N-terminus SCFP3A), pSCYCE(R) (C-terminus of SCFP3A) and pVYNE (N-terminus of Venus) (Gehl et al., 2009) or pESPYNE-Gateway (N-terminus of YFP)/ pESPYCE-Gateway (C-terminus of YFP) (Walter et al., 2004; Schütze et al., 2009).

### Plant protein isolation and detection using immunoblotting

To detect GFP-tagged proteins in *Nicotiana benthamiana,* total protein was extracted from ground leaf material in 2x v/w extraction buffer (8M urea, 100mM Tris pH 6.8, 2% SDS, 10 mM DTT), incubated on ice for 15 min, centrifugated at 13,000g for 20 min at 4°C and then separated on 10% SDS-PAGE, blotted to PVDF membranes and after blocking with 5% milk in PBS detected using monoclonal antibodies (diluted 1:1000) directed against GFP (Chromotec, #029762). The secondary antibody goat anti-rat IgG conjugated to Horseradish peroxidase (ThermoFisher #31470) was used at a dilution of 1:10,000. The proteins were visualized using enhanced chemiluminescence (ECL, home-made recipe). Equal loading of the protein samples was confirmed by Ponceau staining of the blots.

### Protein isolation from yeast and detection using immunoblotting

To obtain a total protein lysate from yeast, the protocol from Yeast Protocols Handbook was followed (Clontech; Yeast Protocols Handbook: www.clontech.com/images/pt/PT3024-1.pdf). Additional details are described in Mazur *et al*. (2017).

### Transient expression of protein in *N. benthamiana* using agro-infiltration

Transient expression of proteins was performed as described by Ma *et al*. (2012). In brief, Agrobacterium GV3101 cells containing the desired constructs were infiltrated into 4-5-week-old *N. benthamiana* leaves (OD_600_=1.0 for each construct). To suppress gene silencing, an *A. tumefaciens* strain GV3101 carrying pBIN61 containing the P19 silencing suppressor of Tomato busy shunt virus (TBSV) (Voinnet et al., 2003), was co-infiltrated with the samples (OD_600_=0.5). Protein accumulation was examined 2-3 days post-infiltration.

### Transient expression of proteins in *N. benthamiana* with anacardic acid treatment

Four-week-old *N. benthamiana* plants were used for agroinfiltration. Three days post-agroinfiltration, 100 μM anacardic acid in 1% DMSO or only 1% DMSO (as a negative control) were directly injected into *N. benthamiana* leaves transiently expressing the BiFC constructs. One and half hours after anacardic acid infiltration, the treated leaf discs were collected and fluorescence was analysed using a Zeiss LSM510 confocal laser microscope. Two independent biological experiments were carried out and a minimum of 50 nuclei for each sample and treatment were observed.

### Transient expression using particle bombardment of Arabidopsis

Complete Arabidopsis rosettes of 4-5-week-old plants grown in 11 hrs light/13 hrs dark (22°C) were placed on 1% agar containing 85 μM benzimidazole (Sigma-Aldrich) and kept in the growth chamber until bombardment. Transformation by particle bombardment was performed as described previously (Schweizer et al., 1999; Shirasu et al., 1999). In brief, 1 μm diameter gold particle were coated with 2.5 μg of each type of plasmid. The PDS1000/HE particle gun with Hepta-adaptor (Bio-Rad) was used according to manufacturer’s protocol using a 900 psi. rupture disk (Bio-Rad). After bombardment, Petri dishes were sealed with medical tape and returned to the growth chamber for 2-3 days before inspection. Plates were kept in the dark from the evening before inspection till loading on the confocal microscopy to retain COP1 in the nucleus.

### GAL4 yeast two-hybrid protein-protein interaction assay

The protocol for yeast two-hybrid was followed as described in (de Folter and Immink, 2011) and further details are described in Mazur *et al.* (2017).

### Confocal microscopy

*N. benthamiana* and Arabidopsis leaves were analyzed 2-3 days post infiltration with *A. tumefaciens* or particle bombardment, respectively. The Hoechst 33342 (Sigma-Aldrich) chromatin stain and FM4-64 (ThermoFisher) membrane dye and endocytic tracer were syringe infiltrated into the leaves at a final concentration of 1 μg/ml and 50 µM in water, respectively, and stored in the dark in a Petri dish on wet paper. FM4-64 could be directly imaged, Hoechst 33342 was incubated for at least an hour before imaging. Accumulation of the tagged proteins was examined in leaf epidermal cells using a Zeiss LSM510 or Nikon A1 confocal laser-scanning microscope. Images at the LSM510 were taken with C-Apochromat 40x water immersive objective with a numerical aperture (NA) of 1.2. At the A1 images were taken with a Plan Fluor 40x oil immersive DIC lens (NA=1.3). SCFP, GFP, chimeric Venus^N^-SCFP^C^, YFP and RFP labelled samples were excited with 458, 488, 488, 514 nm or 568 nm diode lasers. At the LSM510, GFP and the Venus^N^-SCFP^C^ chimera were detected using a 520-555 nm BP filter (BP520-555), SCFP (BiFC) with BP470-500, RFP with BP585-615. At the Nikon A1, SCFP was detected with BP468-502; YFP/GFP with BP500-550, and RFP with BP570-620. At the Nikon A1, for simultaneous Hoechst 33342, SCFP and mRFP imaging, Hoechst 33342 was excited with the 402 nm diode laser and detected between 425-475 nm, SCFP was excited with the 488 nm diode laser and detected between 500-550 nm and mRFP was excited with the 561 nm diode laser and detected between 570-620 nm. FM4-64 was excited with the 514 nm diode laser and detected between 570 and 620 nm. Bright field images were recorded with the transmitted light photomultiplier detector. On both microscopes, co-expressed fluorophores or dyes were excited consecutively to limit bleed through of emission signals between detection channels. For all observations, the pinhole was set at 1 Airy unit. Images were processed using ImageJ (NIH).

## Supplemental Data

**Supplemental Figure S1.** Arabidopsis SUMO1 interacts specifically via a non-covalent SIM-like interaction with SCE1 in Arabidopsis.

**Supplemental Figure S2.** The ternary complex between Arabidopsis SUMO1, SCE1 and SIZ1 is stabilized by the substrate-binding pocket (CAT2) and non-covalent interactions between the different proteins.

**Supplemental Figure S3.** The SUMO1^GG^-SCE1 and SUMO1^GG^-SIZ1 BiFC complex localize to the nucleus in nuclear bodies.

**Supplemental Figure S4.** Alignment of the SCE1 protein sequences of different monocot and dicot plant species showing that SCE1 protein is extremely conserved at the sequence level.

**Supplemental Figure S5.** Co-expression of SCE1·SIZ1 as BiFC pair with an excess of YFP-SUMO1^GG^ drives an increased pool of the SCE1·SIZ1 BiFC pair to small puncta.

**Supplemental Figure S6.** SUMO chain formation accelerates localization of the SUMO1·SCE1 BiFC pair in nuclear bodies.

**Supplemental Figure S7.** Quantification of the co-localization of the SUMO1·SCE1 BiFC pair with COP1.

**Supplemental Figure S8.** Neither mRFP-SCE1 nor SIZ1-mRFP are recruited to CRY2 NBs.

**Supplemental Table S1** SUMO1, COP1, SIZ1 and SCE1 mutants here used.

**Supplemental Table S2** Sequences of oligonucleotides.

**Supplemental Table S3** Clone information.

## Acknowledgments

We kindly acknowledge L. Tikovsky and H. Lemereis for horticultural assistance, the van Leeuwenhoek Centre for Advanced Microscopy, Section Molecular Cytology (SILS-University of Amsterdam) for use of their microscopy facilities, V. Hammoudi for SIZ1*^SP-RING^*. We thank D. Baulcome (University of Cambridge, UK), Dikic (Goethe University, Frankfurt am main, Germany), Dongqing Xu (Peking University, Beijing), J. Haas (Ludwig-Maximilians-University of Munich, Germany), Kudla (WWU Muenster, Germany), Klaus Harter (ZMBP Tübingen, Germany), T. Nagakawa (Shimane University, Japan), Seong Wook Yang (University of Copenhagen, Denmark), Jin Bo Lin (Chinese Academy of Sciences, Beijing, China), Dae-Jin Yun (Gyeongsang National University, Korea) and Ute Höcker (Universität zu Köln) for sharing plasmids and seeds. We thank the ABRC for sending us the CRY1 and CRY2 cDNA clones. The Netherlands Scientific Organisation (ALW-VIDI grant 864.10.004 to HvdB) and the Topsector T&U program Better Plants for Demands (grant 1409-036 to HvdB), including the partnering breeding companies, supported this work; FM is financially supported by Keygene N.V. (The Netherlands).

Author contributions
HB conceptualized the project. MM, MK, FM and HB designed experiments. MM, MK, MAM, FM and RK performed experiments. MM, MK, MA, FM and HB analysed the data. MM, MK and HB wrote the MS. MM, MK, FM, MAM, MP and HB reviewed and edited the MS. HB and MP acquired funding and supervised the project. MM and MK contributed equally to this work.

